# Systematic profiling of growth interactions in human gut microbiome species

**DOI:** 10.64898/2025.12.19.693361

**Authors:** Bolor Buyanbadrakh, Elisabeth Baland, Paula Lambeck, Lucía Pérez Jiménez, Sandra M. Holmberg, Fabiola Puértolas-Balint, Eric Toh, Sun Nyunt Wai, Björn O. Schröder, Madeleine Ramstedt, Shaochun Zhu, André Mateus

## Abstract

Microbial interactions shape the composition and stability of the human gut microbiome. Yet, the molecular mechanisms underlying these relationships remain poorly understood. Here, we systematically profiled 1,224 pairwise growth interactions among 36 representative gut bacterial strains using a spent medium assay. Most interactions were inhibitory and largely explained by resource depletion or medium acidification. We investigated further the molecular basis of a positive interaction, showing that *Clostridium perfringens* promotes the growth of *Mediterraneibacter gnavus* through extracellular vesicles. Additionally, we identified *Veillonella parvula* as a key species capable of modulating environmental pH, thereby enabling the growth of *Parabacteroides merdae*, a strain highly sensitive to acidic conditions. This pH-buffering effect was enhanced by guanine supplementation and persisted in multi-species communities containing different pH-lowering strains from diverse bacterial phyla. This demonstrates the ecological relevance of *V. parvula*, a species that, despite being commonly present in human gut microbiomes, is generally found at low levels. Overall, the comprehensive dataset and mechanistic insights presented here provide a foundation for rational strategies to manipulate microbiome composition and function.

## Introduction

Microorganisms commonly live in complex communities, whose composition is shaped by molecular interactions among the microorganisms that compose them^1,2^. Species may compete for nutrients, secrete inhibitory molecules either into the environment or directly into neighboring cells, or alter the chemical environment (for example, by lowering the pH) to restrict the growth of other species. On the other hand, microorganisms can secrete metabolic by-products that promote the growth of other species. The combined effect of all the interactions results in a community that has a stable composition, unless challenged by strong perturbations such as antibiotic treatment or dietary changes^3–5^. In the context of the human microbiome, this community resilience is critical to prevent colonization by pathogenic species^6–8^.

Studying species interactions is thus key to predict community composition, and ultimately its molecular functions. This can be accomplished by growing one species on sterilized medium in which another species was previously grown (spent medium) ^9–11^, or co-culturing pairs of species together and deconvoluting their individual growth, e.g., by measuring the abundance of species-specific DNA sequences^12–15^ or proteins^16^, or by physically separating species while allowing molecular exchange^17^. These pairwise interactions are generally able to predict bacterial behavior in a community^12,14^, but higher order interactions can also be studied directly by assembling communities of different complexity^18–20^. Alternatively, it is also possible to use computational approaches, such as models based on the metabolic capacity encoded on the genome of each species^21^.

In this work, we used a spent medium approach to systematically profile pairwise growth interactions of 36 representative bacterial strains of the human gut microbiome (**Table S1**). These species cover a large phylogenetic diversity (spanning six major phyla) and exhibit varied abundance and prevalence across individuals^22,23^. We used these data to prioritize the study of molecular mechanisms behind specific interactions, which enables establishing rational strategies to manipulate microbial community composition and function.

## Results

### Binary growth interactions among human gut microbes

To study directional growth interactions among human gut bacteria, we used a spent media assay. First, we collected the spent medium of 36 bacterial strains grown for 24 h in a rich medium (Gifu anaerobic medium; GAM). Then, each individual strain was cultured in each of these spent media or nutrient replenished spent media (spent media mixed with an equal volume of 2x concentrated rich medium). As a control, we grew the same strains in rich medium (**Figure 1a**). Growth was monitored by continuously measuring optical density at 600 nm (OD_600_) for 48 h. Growth dynamics were quantified by calculating the area under the growth curve (AUC; **Table S2**). Due to inconsistent day-to-day variation in lag phase, *Segatella copri* and *Odoribacter splachnicus* were excluded from further growth analyses. However, we still assessed how other strains grew on their spent media, because the yield after 24 h was consistent. In total, this resulted in 1,224 binary interactions being tested (**Figure 1b**; **Table S2**). Growth measurements were highly reproducible, with a median coefficient of variation of 8.5% (interquartile range: 13.6%) for the AUC across replicates (**Figure 1c**).

**Figure 1.**
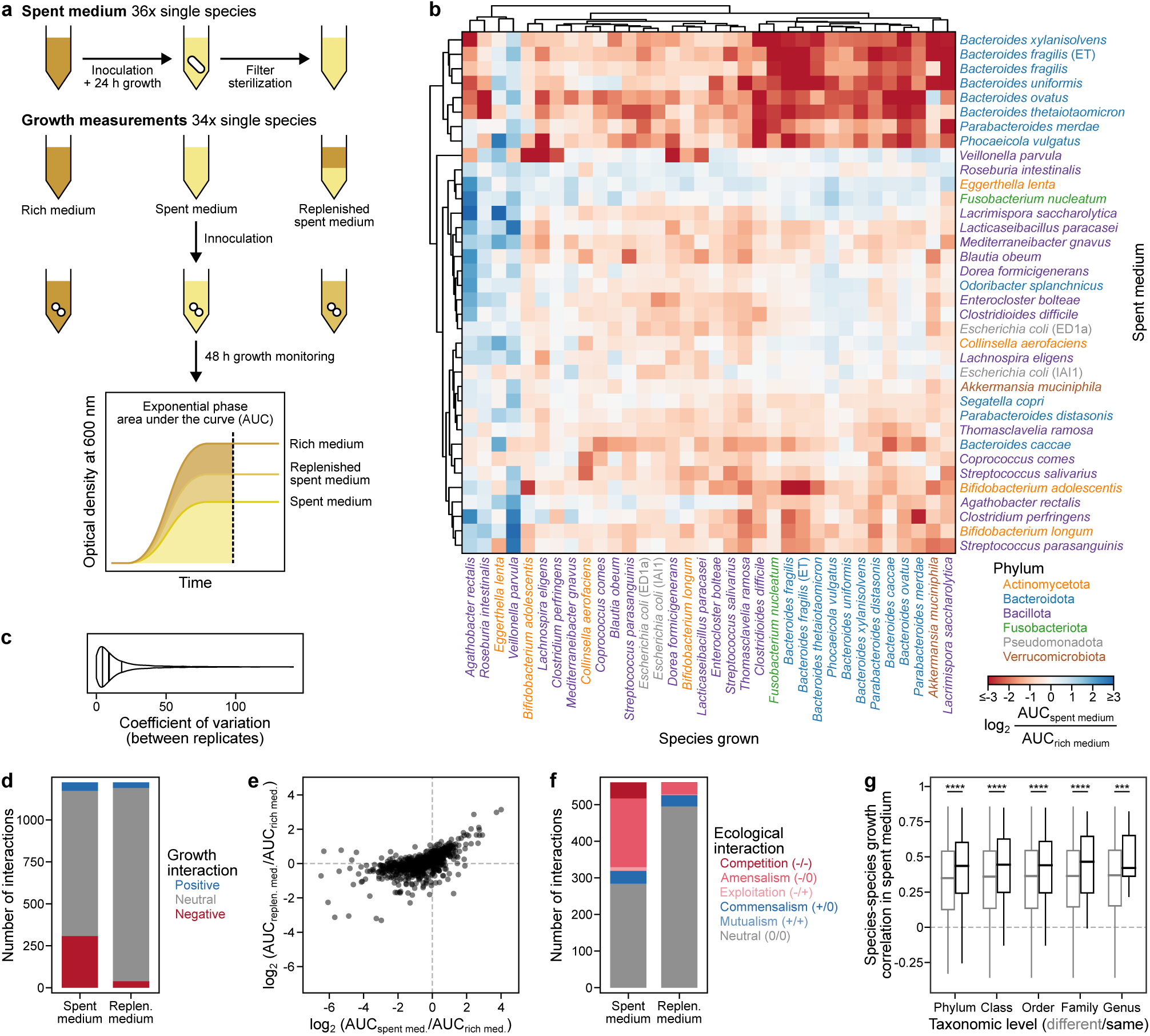
Widespread growth inhibiting and growth promoting ecological interactions among human gut microbes. **(a)** Interactions were measured by spent medium experiments in which one of 36 bacterial strains was grown for 24 h in rich medium followed by filter sterilization. A second strain was then grown in this spent medium, replenished medium, or rich medium as a control and growth was followed by measurements of optical density at 600 nm (OD_600_). The area under the curve of the early stationary phase (AUC) was used as the final growth measurement. **(b)** Pairwise changes in AUC of strains grown (column) in spent medium (row) of another strain compared to rich medium. Strains are color coded according to the phylum to which they belong. **(c)** Reproducibility of growth measurements as measured by the coefficient of variation of the AUC across replicates. Median is highlighted with a thicker line and quartiles with thinner lines. **(d)** Number of interactions with a change of at least 2-fold in spent medium or replenished medium compared to rich medium. **(e)** Comparison of spent medium and replenished medium experiments. **(f)** Number of interactions between two strains using an ecological classification to identify reciprocity. **(g)** Distribution of Pearson correlation coefficients between the growth behavior of pairs of strains (column-wise correlation of values in panel **b**) if they belong or not to the same taxonomic level. Box represents the interquartile range, line represents the median, and whiskers represent the 5^th^ and 95^th^ percentiles. \*\*\**p*<0.001, \*\*\*\**p*<0.0001.

When considering a two-fold change compared to rich medium (|log_2_ fold change|>1), we observed 307 inhibitory interactions (25.1%), and 51 interactions where growth was promoted (4.2%) (**Figure 1d**). The number of inhibitory interactions decreased by more than 8-fold to 38 interactions (3.1%) when spent medium was replenished with nutrients, with most inhibitory effects becoming neutral (**Figure 1e**). This suggests that most inhibitory interactions are due to depleted resources or changes to the media that limit growth (e.g., pH; see below). In contrast, a large proportion of growth promotion interactions (27 out of 58 interactions) were unchanged even after nutrient replenishment, suggesting potential cross-feeding.

We next explored if inhibitory interactions were mostly reciprocal, classified as competition (-/-) in ecology, or if one strain showed exploitation (-/+) or amensalism (-/0). Our results showed that most interactions pointed to amensalism (33.5%), followed by competition (7.8%), with exploitation accounting only for 1.8% of the interactions (**Figure 1f**). Only a small fraction of commensalism (+/0; 6.3%) and mutualism (+/+;0.36%) was observed. It should be noted that these relationships are inferred, since strains were not co-cultured. However, they reflect known interactions, for example, we observed that the spent media of *Fusobacterium nucleatum*, *Streptococcus salivarius*, and *S. parasanguinis* promoted the growth of *Veillonella parvula* (log□ fold-change of 1.4-2.7 compared to rich medium; **Table S2**), which aligns with previous reports showing that *Veillonella* frequently co-occurs with *F. nucleatum* and *Streptococcus* species in oral microbiomes and exhibits strong intergeneric coaggregation^24,25^.

Finally, we noticed that strains that were phylogenetically more closely related tended to behave similarly across the different spent media as indicated by a stronger correlation in their responses to spent media compared to pairs without taxonomic match (**Figure 1g**), with the effect being stronger the more taxonomically related two strains were. This indicates that taxonomic similarity may be a contributing factor to how species interact with, and respond to, the metabolic environment shaped by others.

In summary, we generated a comprehensive dataset of pairwise strain interactions across human gut microbiome bacteria. This highlighted a high prevalence of competition due to resource depletion or changes to the ecosystem, but also some cooperative interactions that are important in shaping community composition.

### *Clostridium perfringens* vesicles enhance the growth of *Mediterraneibacter gnavus*

*Mediterraneibacter gnavus* (formerly *Ruminococcus gnavus*) exhibited enhanced growth when cultured in *Clostridium perfringens* spent medium compared to standard rich medium (**Figure 2a**). To investigate the molecular basis of this effect, we performed a quantitative proteomic analysis of *M. gnavus* grown in *C. perfringens* spent medium versus rich medium. The proteomic profile revealed a decreased abundance of proteins involved in *de novo* nucleotide synthesis, and a concomitant increase in proteins associated with nucleotide salvage pathways (**Figure 2b; Table S3**). This suggested that *M. gnavus* relies less on the synthesis of nucleotides when grown in *C. perfringens* spent medium. Thus, we tested whether nucleic acid components could promote the growth of *M. gnavus* by culturing it in media supplemented with nucleobases, nucleosides, and nucleic acids. However, none of these supplements showed an effect (**Figure 2c; Table S4**), suggesting that a more complex or alternative uptake mechanism may be involved.

**Figure 2.**
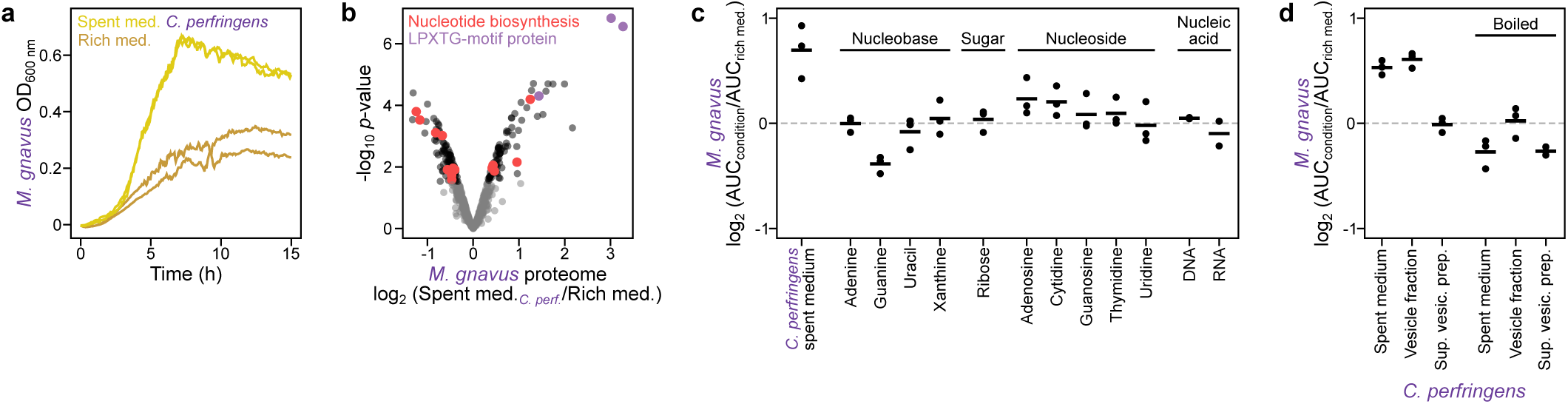
Mechanism of growth enhancement of *Mediterraneibacter gnavus* by *Clostridium perfringens* spent medium. **(a)** Growth over time as measured by OD_600_ of *M. gnavus* in rich medium or *C. perfringens* spent medium. **(b)** Proteome abundance changes of *M. gnavus* grown in *C. perfringens* spent medium compared to rich medium. **(c)** Effect of nucleobases, sugar, nucleosides, and nucleic acids on the growth of *M. gnavus* compared to rich medium. **(d)** Effect of vesicle fraction present in *C. perfringens* spent medium, or the supernatant of the purification, on the growth of *M. gnavus* compared to rich medium. All conditions were also boiled prior to measurement.

Previous studies have reported that *C. perfringens* secretes membrane vesicles containing proteins, peptidoglycan, and nucleic acids^26–28^. The proteome analysis of *M. gnavus* also revealed a strong increase in the abundance of three LPXTG-motif cell wall anchor domain proteins (**Figure 2b**). Despite having an uncharacterized function, these proteins are generally associated with interactions with the environment (e.g., adhesion or colonization) ^29,30^. Using dynamic light scattering, we observed particles with an average diameter of 71 nm in *C. perfringens* spent medium, while rich medium contained particles with an average diameter of 149 nm (**Figure S1**). We then isolated the crude extracellular vesicle fraction from the spent medium of *C. perfringens* by ultracentrifugation, and confirmed the presence of particles with an average size of 58 nm (**Figure S1**). *M. gnavus* exhibited increased growth when cultured in rich medium supplemented with vesicles isolated from *C. perfringens*, whereas no enhancement was observed in rich medium lacking vesicle supplementation. (**Figure 2d**; **Table S5**). Importantly, growth was not promoted when *M. gnavus* was grown in the supernatant collected after ultracentrifugation (**Figure 2d**), which lacked detectable particles when analyzed by dynamic light scattering (**Figure S1**). Furthermore, heat treatment of the vesicle fraction abolished the growth-promoting effect, indicating that the active components are heat-labile.

In summary, *C. perfringens* promotes the growth of *M. gnavus* by releasing extracellular vesicles. As previously reported^26–28^, these vesicles contain nucleic acid components that may contribute to *M. gnavus* metabolism, and our data further suggest that heat-labile factors and/or intact vesicle structure are necessary for the observed growth enhancement.

### pH changes in the medium are crucial mediators of inter-microbial interactions

Since bacteria can modify and are affected by environmental pH, we studied how pH changes in the spent medium of one strain affected the growth of the cultivated strain. Overall, we observed a positive correlation between the pH of spent media (**Table S1**) and the log_2_ fold change of growth in spent media compared to rich medium (*r* = 0.49, *p* = 10^-73^; across all strain-spent media combinations), indicating that lower pH generally results in reduced growth. This correlation was particularly strong in the Bacteroidota, Fusobacteriota and Pseudomonadota phyla (*r* >0.5), with the former two being more affected at lower pH (**Figure 3a**). In contrast, the Actinomycetota phylum and some strains of the Bacillota phylum (e.g., *V. parvula*) showed weak or negative correlations, implying potential acid tolerance or buffering capacities.

**Figure 3.**
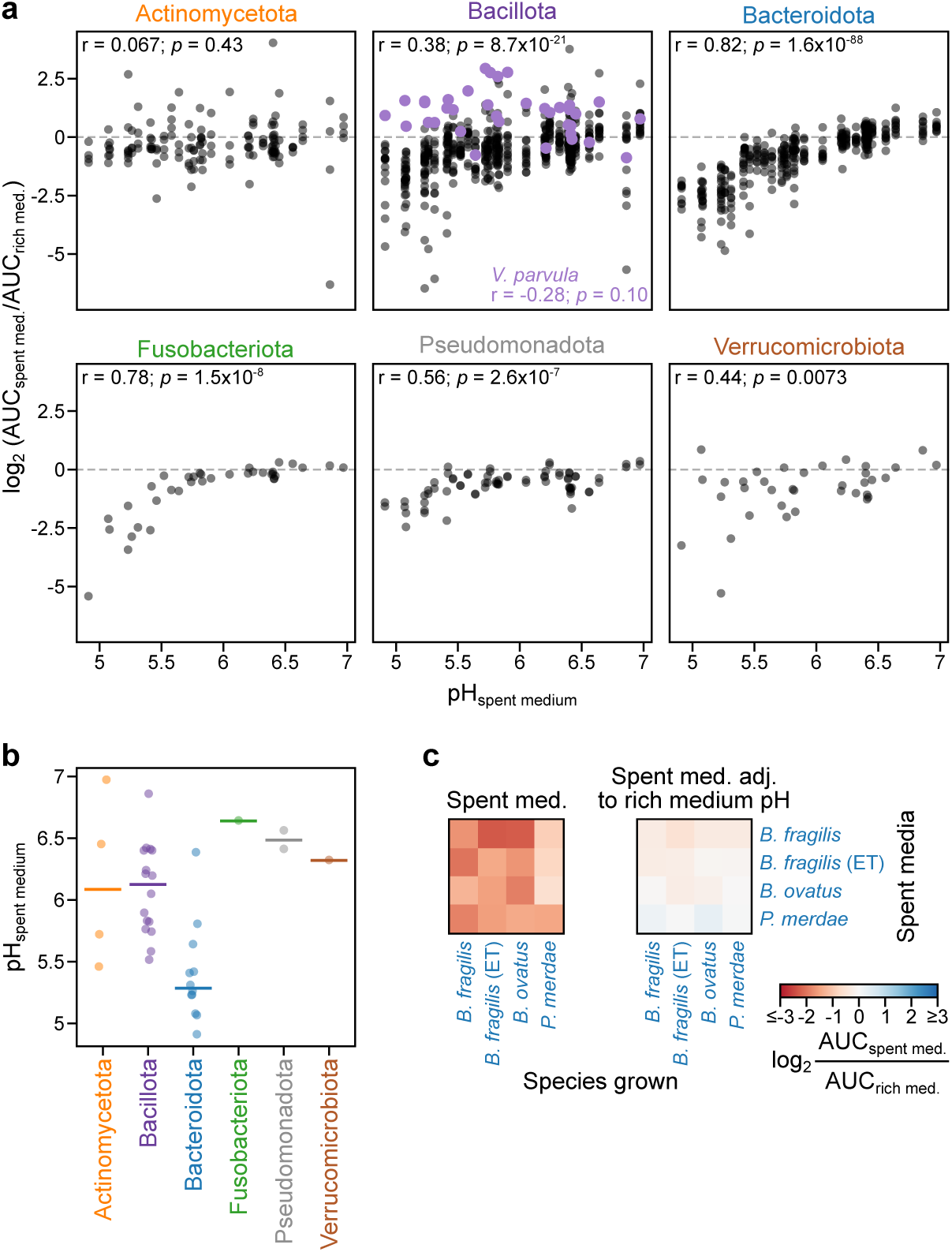
Negative growth interactions are largely explained by lower environmental pH. **(a)** Comparison of changes in growth in spent medium with the pH of the spent medium separated by the phylum of the strain that was grown. **(b)** pH of spent medium of all strains tested in this study according to the phylum to which they belong. **(c)** Pairwise changes in AUC of strains grown (column) in spent medium (row) of another strain compared to rich medium for selected Bacteroidota strains that show lower growth in acidic pH. Adjustment of the pH of spent medium to that of rich medium reduces the growth inhibitory effect.

We thus wondered if growth inhibition between Bacteroidota phylum strains was due to limited nutrient availability or due to the low pH of their spent media, which consistently exhibited the lowest pH values among all tested strains (**Figure 3b**). Of note, replenishing the spent medium restored growth to similar levels as rich medium, but also increased the pH (**Table S1**). To directly assess the effect of pH, we measured the growth of *B. ovatus*, *B. fragilis*, enterotoxin-producing *B. fragilis* (*B. fragilis* ET), and *P. merdae* in both their own and each other’s spent media before and after adjusting the pH to that of rich medium. Upon pH adjustment of the spent medium, the previously observed growth inhibition was reversed (**Figure 3c**; **Table S6**), indicating that low pH is the primary driver of growth inhibition, rather than nutrient exhaustion.

Overall, we observed that media acidification limits the growth of multiple strains. Although the degree of inhibition is strain-specific, some phyla both induce stronger pH reductions and exhibit greater sensitivity to acidic conditions.

### *V. parvula* possesses pH-neutralizing capabilities

*V. parvula* showed a unique behavior compared to other bacteria, since its growth was promoted by 53% of the other strains (**Figure 1b**). Further, it experienced the strongest growth stimulation when cultivated in the spent medium from *Bifidobacterium longum*, *Lacticaseibacillus paracasei*, *Streptococcus parasanguinis*, and *Clostridium perfringens*, all of which slightly acidified the pH of the medium to approximately 5.8 (**Table S1**). This prompted us to examine if *V. parvula* possessed pH-neutralizing capabilities. To study this, we grew *V. parvula* in rich medium adjusted to varying initial pH, and measured the pH and overall growth after 48 hours of incubation. As a comparison, we included *Parabacteroides merdae*, which displayed impaired growth at low pH and acidified the medium. The results showed that *V. parvula* increased the pH of the medium, with the strongest activity observed at pH 5 (**Figure 4a**; **Table S7**). Below pH 5, there was no bacterial growth. This pH-increasing effect gradually diminished as the initial pH increased from 5 to 7, indicating that its ability to raise pH is most pronounced under acidic stress.

**Figure 4.**
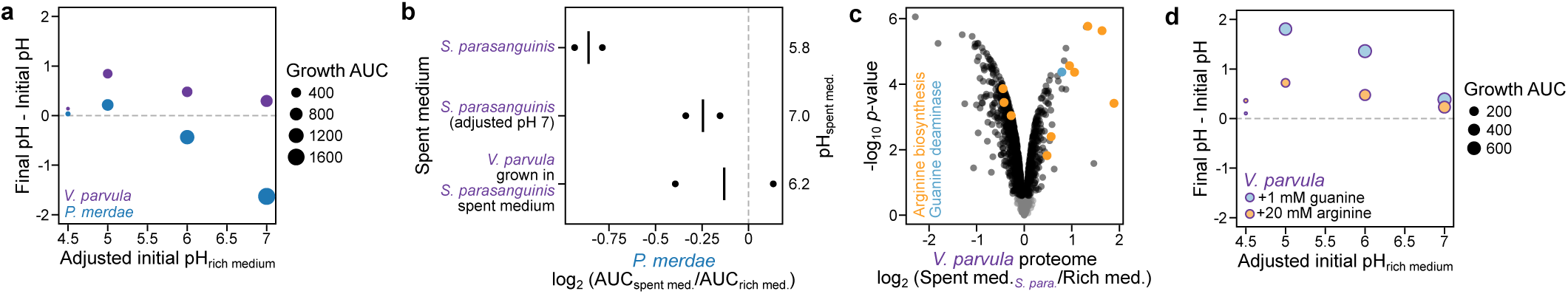
*Veillonella parvula* increases environmental pH and promotes the growth of low pH-sensitive strains. **(a)** Changes in pH of medium induced by the growth of *V. parvula* or *P. merdae* depending on the initial pH of rich medium. Size of points corresponds to the growth AUC in that condition and highlights that the lack of change in pH at the lowest pH is due to the absence of growth. **(b)** Growth of *P. merdae* when grown in *S. parasanguinis* spent medium, this spent medium adjusted to pH 7, or in the spent medium of *V. parvula* grown on *S. parasanguinis* spent medium. All conditions are compared to rich medium (two individual replicates and the average are shown), and initial pH of spent medium is highlighted on the right. **(c)** Proteome abundance changes of *V. parvula* grown in *S. parasanguinis* spent medium compared to rich medium. **(d)** As in panel **a**, but with the addition of 1 mM guanine or 20 mM arginine prior to adjusting the pH of rich medium.

To test whether *V. parvula* could modify acidic spent media and make it more favorable for pH-sensitive strains, we incubated *P. merdae* (a pH-sensitive strain) in *S. parasanguinis* spent medium, which has a pH of 5.8. As expected, the growth of *P. merdae* was inhibited in *S. parasanguinis* spent medium when compared to rich medium, with this effect being alleviated when the pH of this spent medium was adjusted to pH 7 (**Figure 4b**; **Table S8**). Similarly, when *V. parvula* was grown in the spent medium of *S. parasanguinis*, the pH increased modestly to 6.2, and *P. merdae* growth was improved. These results demonstrate that *V. parvula* can raise the pH of the media, thereby reducing acid-driven growth suppression of sensitive strains.

To investigate the molecular mechanisms underlying this pH increase, we quantified the proteome of *V. parvula* grown in *S. parasanguinis* spent medium and in rich medium. This analysis showed an increased abundance of proteins involved in arginine biosynthesis, guanine deaminase (an enzyme that converts guanine to xanthine), and xanthine permease in the spent medium condition compared to rich medium (**Figure 4c**; **Table S9**). Therefore, we tested if the addition of arginine or guanine to rich medium adjusted to different pH would lead to increased medium pH. While the addition of guanine led to a stronger increase in pH (**Figure 4d; Table S7**), the addition of arginine led to an effect similar to rich medium alone (compare **Figure 4d** with **Figure 4a**). It is important to note that pH was adjusted after the addition of guanine and arginine, potentially masking the pH increasing effects of these molecules.

In summary, *V. parvula* increased the pH of the extracellular environment, and this effect was exacerbated by guanine. This pH-neutralizing property can rescue the growth of species that are particularly affected by low pH (e.g., Bacteroidota).

### *V. parvula* pH-neutralizing effect plays an important role in community composition

We pondered if the capacity of *V. parvula* to neutralize the pH of the medium would play a role in a community setting. Thus, we assembled three-member communities consisting of *V. parvula, P. merdae* (a pH-sensitive strain), and an acidifier strain (**Figure 5a**). We chose four phylogenetically diverse acidifying strains: *S. parasanguinis* and *Lacticaseibacillus paracasei* from the Bacillota phylum, *Bifidobacterium adolescentis* from the Actinomycetota phylum, and *Bacteroides ovatus* from the Bacteroidota phylum. In all co-cultures with acidifying strains, *P. merdae* exhibited reduced growth and a lower pH compared to its monoculture (**Figure 5b-c**; **Figure S2**; **Table S10**). *B. ovatus* growth rate was slightly higher than *P. merdae* (**Figure S2e** and **g**), resulting in a stronger decrease in pH than the other acidifying strains (**Figure S2f**), both of which prevented *P. merdae* from growing in co-culture. The addition of *V. parvula* to these communities consistently enhanced *P. merdae* growth and elevated the pH (**Figure 5b**; **Figure S2**), demonstrating capacity of *V. parvula* to increase the environmental pH in diverse community settings and promote the growth of low pH-sensitive strains.

**Figure 5.**
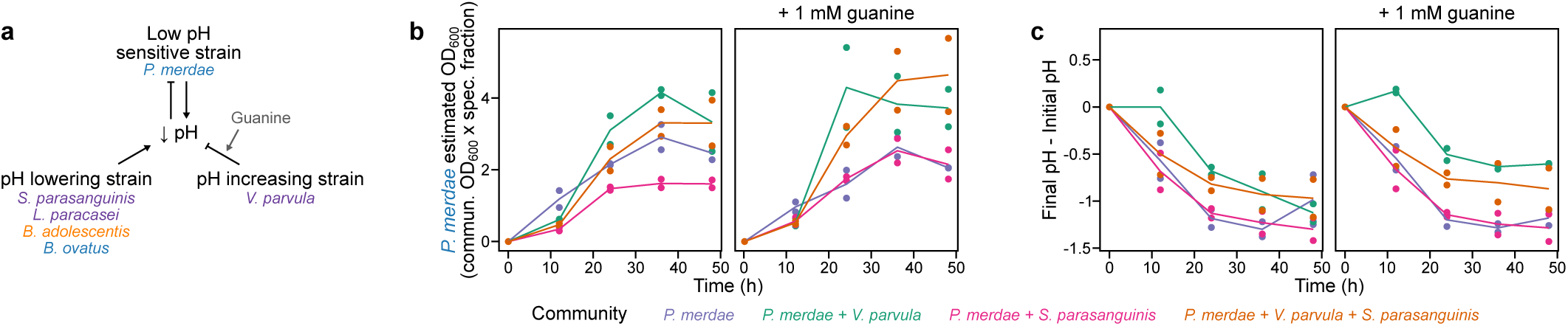
*Veillonella parvula* promotes the growth of a pH-sensitive strain in a community with a pH-lowering strain. **(a)** Model of contributing effects of each strain to the community. **(b)** *P. merdae* growth in different communities over time without or with the addition of guanine. Growth was calculated by multiplying the OD_600_ of the community by the fractional abundance of *P. merdae* in the community as determined by qPCR. **(c)** Environmental pH of different cultures over time.

Since we identified that guanine strengthens the pH-increasing capacity of *V. parvula*, we evaluated its role in these communities (**Figure 5a**). As expected, the addition of guanine promoted *P. merdae* growth and increased the pH in communities in which *V. parvula* was present (**Figure 5b**; **Figure S2**) except for *B. ovatus* communities due to the aforementioned reasons.

In conclusion, *V. parvula* supports the growth of strains sensitive to low pH in a community, with this effect being amplified by the addition of guanine.

## Discussion

We systematically measured all mutual growth interactions between a group of representative strains of the human gut microbiome. For scalability, the analysis was limited to pairwise interactions, since testing all possible community compositions of our group of 36 strains would require testing over 10^10^ conditions^19^. Pairwise interactions generally enable predicting the overall composition of a community ^12,14^. However, emergent behaviors of higher order interactions are possible^31^, as evidenced by our communities with *P. merdae* and *B. ovatus*, and warrant further study. We used a spent medium approach, which disregards contact-dependent mechanisms^32^, but facilitates growth measurements. It also eases mechanistic follow-up work, particularly when using an approach based on proteomics, due to the high dynamic range of the proteome and the general dominance of one strain in co-cultures^16^, or metabolomics, due to the difficulty in identifying which strain consumed or produced a metabolite^14^. In this work, all experiments were performed in rich medium, which neglect specific nutrients that are present in the gut (e.g., complex carbohydrates), and interactions with the host (both the environment it provides, and how it reacts to the presence of microbes). Yet, our approach is applicable to other media and could enable the discovery of further interaction mechanisms in the future.

To quantify interspecies effects, we employed a simple metric that integrates growth rate and overall biomass yield (area under the curve). However, the provided time-resolved growth measurements for all experiments enable the extraction of growth parameters. These can be integrated with computational approaches to predict community dynamics more accurately. Incorporating metabolic modeling^21,33^ may further illuminate the mechanistic basis of the observed interactions.

Our results highlighted a general prevalence of negative interactions, in accordance with previous studies^2,34^. Strains tended to grow more poorly on spent media derived from closely related taxa, likely reflecting competition for similar metabolic niches. Yet, we observed that at least for selected Bacteroidota strains, the low pH of the spent medium was sufficient to limit growth, and this is likely to explain other interactions^10–12,35,36^.

The specific interactions identified in our screening approach are in line with effects that have previously been observed *in vivo*. For example, we detected a strong inhibitory effect of *Clostridioides difficile* spent medium across multiple strains (32% of the strains showed a 1.5-fold lower growth in *C. difficile* spent medium compared to rich medium), in agreement with the role of this opportunistic pathogen in decreasing microbiome diversity^37^. We also identified that *M. gnavus* growth can be promoted by vesicles from another species of the Clostridia class. *M. gnavus* is known to exchange DNA with other members of the same species via membrane vesicles^38^, it is possible that this extends to other related species. Interestingly, *M. gnavus* produces an anti-*C. perfringens* bacteriocin^39,40^, emphasizing strong growth dynamics between these species.

Our study also recapitulated the interactions of *Veillonella parvula* with lactate-producing species^41^, and its capacity to increase environmental pH. This increase facilitated the growth of low pH-sensitive strains, which highlights the role in communities for this prevalent (>50% of human gut microbiomes contain this species^22,23^), but low abundant species (generally <1% abundance in gut microbiomes^22,23^, and 2-8% in our simple communities). The pH-increasing capacity of this species was enhanced by the addition of guanine through an as-yet unidentified mechanism, but which could be related to the deamination of this base. The upregulation of the arginine pathway when *V. parvula* was grown in *S. parasanguinis* spent medium could indicate a need for the synthesis of this amino acid and not be involved in a pH-increasing mechanism, since *S. parasanguinis* is strongly arginolytic^25,42^.

In summary, we provide a comprehensive dataset of interactions among human-relevant gut bacteria. Our approach to suggest interaction mechanisms at the molecular level, by measuring proteome changes of one strain in spent medium of another compared to rich medium, can be applied to other identified interactions. Understanding the molecular mechanisms behind interactions can enable strategies for the rational manipulation of gut microbial communities.

## Methodology

### Spent medium assay

Spent media was prepared fresh by growing each strain in Gifu Anaerobic Medium (GAM) under anaerobic conditions at 37□ °C for 24□ h inside a Coy anaerobic chamber (5% H_2_, 10% CO_2_, 85% N_2_). Cultures were centrifuged at 2000□ × □g for 30 □min at 4 □°C, and the supernatant was filtered through a 0.2 □µm membrane filter (Filtropur S, Sarstedt) to remove cells. The resulting cell-free supernatant (spent medium) was used immediately for growth assays. For growth assays, 150□ µL of medium either standard GAM, spent medium, or a replenished medium composed of 50% spent medium and 50% 2x concentrated GAM was added to each well of a 96-well plate. To this, 50 □µL of a diluted overnight bacterial culture was added, resulting in a final OD_600_ of 0.01. Each condition was tested at least in duplicate. Plates were sealed with a breathable membrane (Breathe-Easy, Diversified Biotech) and incubated at 37□ °C in a Coy anaerobic chamber, and OD_600_ was recorded every 3 minutes for 48 hours using a Cerillo Stratus plate reader.

### Growth assay data analysis

Growth curves were first trimmed at strain-specific time points when they reached stationary phase to avoid diluting growth responses the same time point was used for all experiments of the same strain (**Table S1**). All growth curves were then visually inspected, and replicates where the rich medium condition deviated from the overall behavior of that strain in rich medium (observed in all other experiments) were removed in those cases all conditions of that replicate (rich medium, spent medium, and replenished media) were removed. The area under the growth curve (AUC) was then calculated using the trapezoid method and fold changes to the rich medium condition were calculated. In rare cases in which the AUC was negative in spent or replenished media, due to lack of growth or growth below the detection limit of the plate reader, the AUC was set to the lowest AUC value for that strain in the respective media (so that a fold-change could be calculated). For all follow-up experiments, the spent medium assay was repeated in that specific condition.

### Proteomics analysis

Bacterial cells were harvested from a 24 h culture by centrifugation at 3000 x g for 15 min at 4 □°C. The pellet was resuspended in PBS containing 2% SDS and incubated at 98□ °C for 10 min. Protein concentration was determined using the BCA assay. 6 µg of each sample was denatured with a final concentration of 2% SDS and 20mM TCEP. Samples were digested with a modified sp3 protocol^43^, as previously described^44^. Briefly, samples were added to a bead suspension (10 μg of beads; Sera-Mag Speed Beads, 4515-2105-050250, 6515-2105-050250, in 10 μl 15% formic acid and 30μl ethanol) and incubated shaking for 15 min at room temperature. Beads were then washed four times with 70% ethanol. Proteins were digested overnight by adding 40 μl of 5 mM chloroacetamide, 1.25 mM TCEP, and 200 ng trypsin in 100 mM HEPES pH 8.5. Peptides were eluted from the beads and dried under vacuum prior to labeling with tandem mass tags (Thermo Fisher Scientific). Labeled peptides were pooled and desalted with solid-phase extraction using a Waters OASIS HLB μElution Plate (30 μm). Samples were fractionated into 48 fractions on a reversed-phase C18 system running under high pH conditions, with every sixth fraction being pooled together. Samples were analyzed by LC-MS/MS, using a data-dependent acquisition strategy on a Thermo Fisher Scientific Vanquish Neo LC coupled with a Thermo Fisher Scientific Orbitrap Exploris 480. Raw files were processed with MSFragger^45^ against the uniprot proteome of the grown and spent medium strains (UP000000818 and UP000004410 for the *M. gnavus* grown in *C. perfringens* spent medium experiment, and UP000778864 and UP000001502 for the *V. parvula* in *S. parasanguinis* spent medium experiment) using standard settings for TMT. The median abundance in each TMT channel was normalized to the median intensity of all TMT channels and statistical significance was determined using limma^46^.

### Growth assays for measuring the effect of nucleotides on *M. gnavus*

Adenine, guanine, uracil, and xanthine were dissolved in 0.1 □M NaOH and added to rich medium at a final concentration of 1□ mM. The addition of NaOH did not significantly alter the pH; a rich medium containing the same volume of NaOH was used as a control. D-ribose was dissolved in water and added to rich medium at a final concentration of 1□ mM. Adenosine, cytidine, guanosine, 2-deoxythymidine were dissolved in PBS and added to rich medium at a final concentration of 1 □mM. DNA (low molecular weight from salmon sperm) and RNA (from yeast) were dissolved in PBS and added to rich medium at a final concentration of 0.5□ mg/mL. *M. gnavus* was inoculated into each condition at an initial OD_600_ of 0.01, and cultures were incubated anaerobically at 37□ °C. Growth was monitored for 48 h as described above, with data analyzed in a similar manner.

### Crude extracellular vesicle isolation and growth assays

Vesicles were isolated from 500 mL of *Clostridium perfringens* culture grown anaerobically in rich medium for 24 h. Cultures were first centrifuged at 2000 □x □g for 30 minutes at 4□ °C to remove bacterial cells. The resulting supernatant was sterilized by filtration through a 0.2□ µm membrane filter (Thermo Fisher Scientific 0.2 aPES) and subsequently ultracentrifuged at 29,000 □r.p.m. for 4 h at 4°C using a Ti 45 rotor (Beckman Coulter Optima) to pellet crude extracellular vesicles. The pellet was resuspended in 1□ mL PBS and stored at −20°C until further use. To assess the effects of vesicles and related fractions on *M. gnavus* growth, isolated vesicles were diluted 1:50 in GAM prior to testing. *C. perfringens* spent medium (from which the vesicles were isolated), and vesicle-free supernatant (the supernatant after ultracentrifugation) were tested without dilution. Boiled versions of crude vesicles, spent medium, and the vesicle-free supernatant were prepared by incubating each sample at 98°C for 10 min. *M. gnavus* was inoculated into each condition at an initial OD_600_ of 0.01, and cultures were incubated under anaerobic conditions at 37°C. Growth was monitored for 48 h as described above, with data analyzed in a similar manner.

### Dynamic light scattering analysis of extracellular vesicles

The size of extracellular vesicles was determined using dynamic light scattering (DLS) with a Malvern Zetasizer Nano ZS at 25□ °C. The vesicle samples were diluted two-fold in PBS, and all other samples were analyzed without dilution. Three biological replicates of each sample were independently prepared, and each one was measured three consecutive times and averaged.

### Strain growth at different pH

Stocks of guanine (1□mM, prepared by dissolving in 1□ M NaOH) and arginine (20 □mM, prepared by dissolving in sterile water) were prepared. These stocks were first added to one-third of the required volume of 2x GAM by carefully dropping under slow stirring. The pH of GAM, GAM supplemented with guanine, or GAM supplemented with arginine was then adjusted to 4.5, 5, 6, or 7 using 1□ M HCl and filter sterilized. The mixture was transferred to an anaerobic chamber, where two-thirds of 2x GAM was added, and finally diluted to the desired volume with sterile double-distilled water. The final media were incubated in anaerobic chamber at least 12 h before use. *V. parvula* was grown overnight in GAM supplemented in 0.5% lactate to improve growth. *V. parvula* and *P. merdae* were inoculated into each condition at an initial OD_600_ of 0.01, and cultures were incubated under anaerobic conditions at 37°C. Growth was monitored over a 48 h period as described above, with data analyzed in a similar manner.

### Community assembly and growth conditions

Microbial communities consisting of two or three bacterial strains (*P. merdae, V. parvula, S. parasanguinis, L. paracasei, Bifidobacterium adolescentis, and Bacteroides ovatus*) were assembled and cultured under two conditions: GAM and GAM supplemented with 1 mM guanine. Following guanine addition, the pH of the supplemented medium was adjusted to match that of the standard GAM to ensure consistency across conditions. Each strain was inoculated into the respective medium at an initial OD_600_ of 0.01. Cultures were incubated under anaerobic conditions at 37°C, and OD_600_ was measured at 12, 24, 36, and 48 h. At each time point, a volume of sample corresponding to one unit of OD_600_ was collected. Cells were harvested by centrifugation, and the resulting pellets were stored at - 20□ °C for downstream analyses.

### Quantitative polymerase chain reaction (qPCR)

Genomic DNA was extracted from bacterial community cultures using the MP FastDNA™ kit (MP Biomedicals) according to the manufacturer’s instructions. DNA concentration and purity were assessed using a NanoDrop spectrophotometer (Thermo Fisher Scientific). PCR was performed on a Bio-Rad CFX Duet real-time PCR system using 2x iQ™ SYBR® Green Supermix (Bio-Rad). Each 10□ µL reaction contained 1x iQ SYBR Green Supermix, 0.2□ µM of each primer (**Table S11**), and 1□ ng of template DNA, with nuclease-free water added to the final volume. The thermal cycling conditions were: initial denaturation at 95 □°C for 3□ min, followed by 35 cycles of denaturation at 95 □°C for 15□ s, annealing at 59°C for 30 s and extension at 72°C for 30□ s. Absolute gene copy numbers were determined using a standard curve generated from serial 10-fold dilutions of custom-made plasmids containing the target genes (10^2^–10^8^ copies per reaction). Absolute copy numbers were corrected by the gene copy number in the genome. The fraction of each species in the community was calculated as the ratio of the corrected copy number of that species to the summed corrected copy numbers of all species present in the community. OD_600_ for each species was estimated as the total OD_600_ of the community multiplied by the fraction of the species.

### Data availability

Raw growth data can be found at https://zenodo.org/records/17456178 or in the supplementary tables. The mass spectrometry proteomics data have been deposited to the ProteomeXchange Consortium via the PRIDE partner repository with the dataset identifier PXD070029 (during review data can be accessed using username; password: K8qfsTSkSyXn).

### Code availability

The code to reproduce the growth assay data analysis is available at https://github.com/andrenmateus/SpeciesSpeciesInteractions.

## Acknowledgements

This work was supported by the Swedish Research Council (grant number: 2022-02958 to A.M.; 2022-02973 to M.R.; 2022-00981 to S.N.W.), the European Research Council (grant number: 101076015), the Knut and Alice Wallenberg Foundation (Wallenberg Academy Fellows 2023), Kempestiftelserna (grant number: JCK3126), and Jeanssons stilftelser (grant number: J2022-0003). MIMS is supported by the Swedish Research Council (grant number: 2021-06602). We acknowledge the Umeå Hypoxic Research Facility (UHRF) at Umeå university.

## Author contributions

B.B., and A.M. designed the study. B.B., E.B., P.L., and L.P.J. performed growth experiments. B.B., S.H., F.P-B., and B.S. performed and analyzed bacterial community experiments. B.B., E.T., S.N.W. and M.R. performed and analyzed the vesicle experiments. B.B., S.Z., and A.M. performed and analyzed the proteomics experiments. B.B., and A.M. drafted the manuscript, which was reviewed and edited by all authors. A.M. acquired funding and supervised the study.

## Supplementary material

### Supplementary tables

**Table S1. Strains included in this study and pH of spent medium after 24 h growth.**

**Table S2. Species-species growth results.**

**Table S3. Proteome analysis of *Mediterraneibacter gnavus* grown on *Clostridium perfringens* spent medium.**

**Table S4. Effect of *Clostridium perfringens* spent medium, nucleobases, ribose, nucleosides, and nucleic acids on the growth of *Mediterraneibacter gnavus*.**

**Table S5. Effect of *Clostridium perfringens* spent medium and purified membrane vesicles on the growth of *Mediterraneibacter gnavus*.**

**Table S6. Effect of spent medium pH on the growth of Bacteroidota species.**

**Table S7. Effect of pH and addition of guanine and arginine on the growth of *Veillonella parvula* and *Parabacteroides merdae*.**

**Table S8. Effect of *Streptococcus parasanguinis* spent medium pH, and a spent medium pH modifying strain (*Veillonella parvula*) on the growth of *Parabacteroides merdae*.**

**Table S9. Proteome analysis of *Veillonella parvula* grown on *Streptococcus parasanguinis* spent medium.**

**Table S10. Effect of *Veillonella parvula* on bacterial communities.**

**Table S11. Primers used for qPCR.**

### Supplementary figures

**Figure S1.**
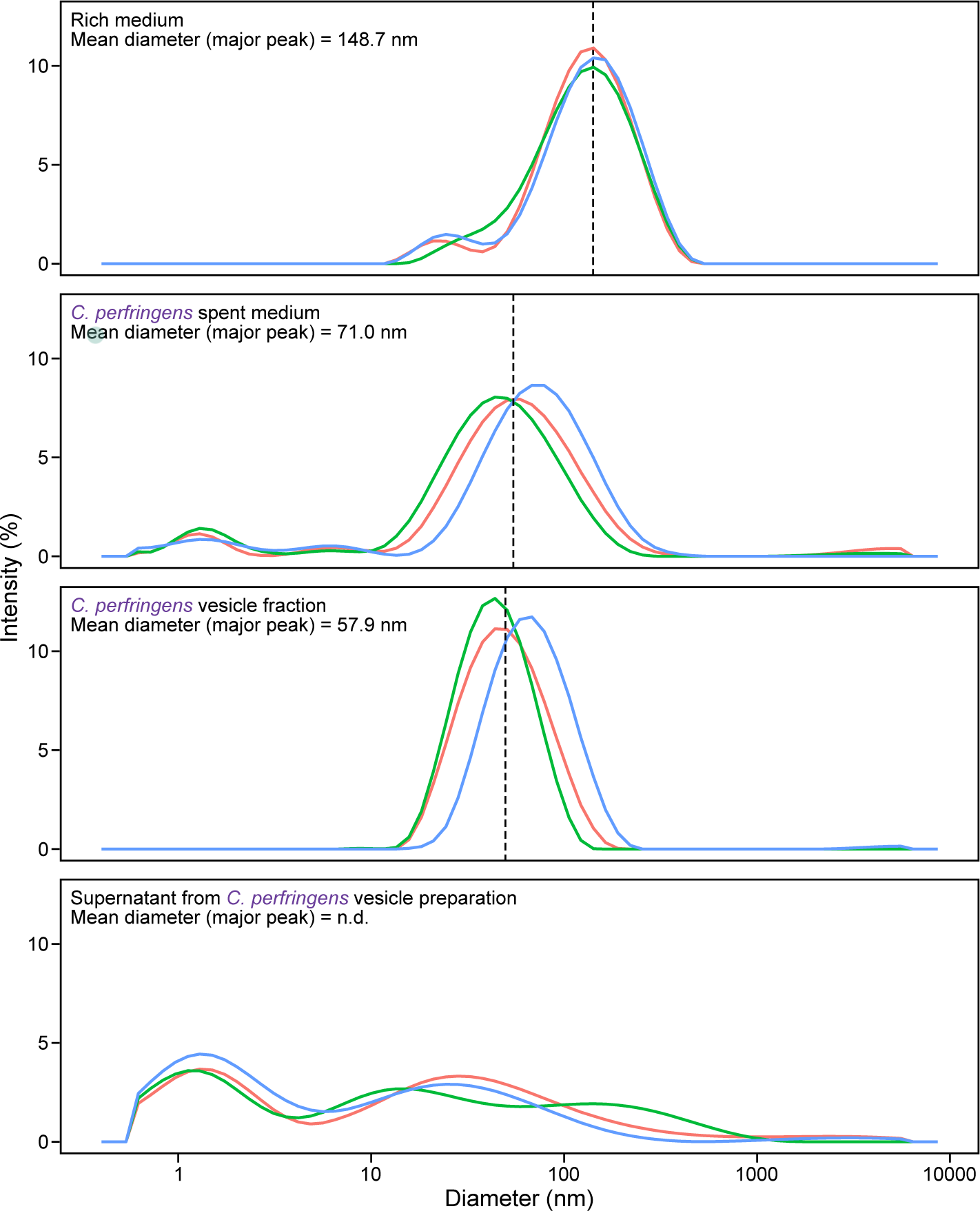
Particle size distribution of rich medium, *C. perfringens* spent medium, vesicles isolated by ultracentrifugation of *C. perfringens* spent medium, and the supernatant after the ultracentrifugation process.

**Figure S2.**
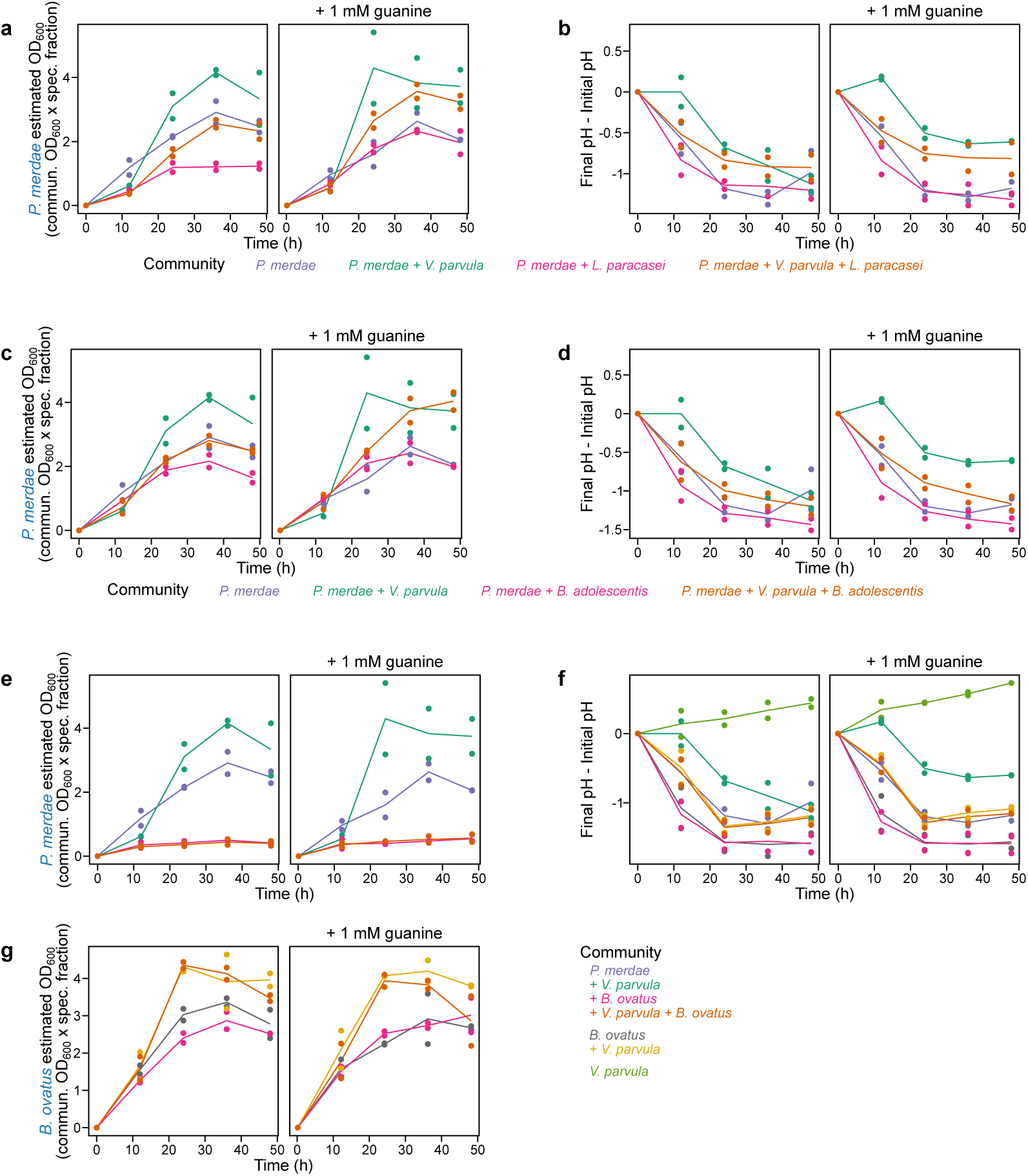
Effect of *V. parvula* on the growth of *P. merdae* and pH in bacterial communities with (a-b) *L. paracasei*, (c-d) *B. adolescentis*, and (e-f) *B. ovatus* (as in Figure 5b). (g) *B. ovatus* growth in the latter community.

## References

1 Zengler, K. & Zaramela, L. S. The social network of microorganisms - how auxotrophies shape complex communities. Nat Rev Microbiol 16, 383–390 (2018). 10.1038/s41579-018-0004-5

2 Kost, C., Patil, K. R., Friedman, J., Garcia, S. L. & Ralser, M. Metabolic exchanges are ubiquitous in natural microbial communities. Nat Microbiol 8, 2244–2252 (2023). 10.1038/s41564-023-01511-x

3 Faith, J. J. et al. The long-term stability of the human gut microbiota. Science 341, 1237439 (2013). 10.1126/science.1237439

4 Caballero-Flores, G., Pickard, J. M. & Nunez, G. Microbiota-mediated colonization resistance: mechanisms and regulation. Nat Rev Microbiol 21, 347–360 (2023). 10.1038/s41579-022-00833-7

5 Butler, S. & O’Dwyer, J. P. Stability criteria for complex microbial communities. Nat Commun 9, 2970 (2018). 10.1038/s41467-018-05308-z

6 Garcia-Santamarina, S. et al. Emergence of community behaviors in the gut microbiota upon drug treatment. Cell 187, 6346–6357 e6320 (2024). 10.1016/j.cell.2024.08.037

7 Griesshammer, A. et al. Non-antibiotics disrupt colonization resistance against enteropathogens. Nature 644, 497–505 (2025). 10.1038/s41586-025-09217-2

8 Kumar, A. et al. Identification of medication-microbiome interactions that affect gut infection. Nature 644, 506–515 (2025). 10.1038/s41586-025-09273-8

9 Ratzke, C., Barrere, J. & Gore, J. Strength of species interactions determines biodiversity and stability in microbial communities. Nature Ecology & Evolution 4, 376–383 (2020). 10.1038/s41559-020-1099-4

10 Weiss, A. S. et al. In vitro interaction network of a synthetic gut bacterial community. ISME J 16, 1095–1109 (2022). 10.1038/s41396-021-01153-z

11 Bonillo-Lopez, L. et al. In vitro metabolic interaction network of a rationally designed nasal microbiota community. iScience 28, 113114 (2025). 10.1016/j.isci.2025.113114

12 Venturelli, O. S. et al. Deciphering microbial interactions in synthetic human gut microbiome communities. Mol Syst Biol 14, e8157 (2018). 10.15252/msb.20178157

13 Friedman, J., Higgins, L. M. & Gore, J. Community structure follows simple assembly rules in microbial microcosms. Nat Ecol Evol 1, 109 (2017). 10.1038/s41559-017-0109

14 Ho, P. Y., Nguyen, T. H., Sanchez, J. M., DeFelice, B. C. & Huang, K. C. Resource competition predicts assembly of gut bacterial communities in vitro. Nat Microbiol 9, 1036–1048 (2024). 10.1038/s41564-024-01625-w

15 van den Berg, N. I., et al. Emergent survival and extinction of species within gut bacterial communities. bioRxiv, 2024.2004.2029.591619 (2024). 10.1101/2024.04.29.591619

16 Kamrad, S. et al. Interspecies interactions drive bacterial proteome reorganisation and emergent metabolism. bioRxiv, 2025.2005.2014.653412 (2025). 10.1101/2025.05.14.653412

17 Zhu, J. et al. Systematic pairwise co-cultures uncover predominant negative interactions among human gut bacteria. Microbiome 13, 161 (2025). 10.1186/s40168-025-02156-0

18 Ishizawa, H., Tashiro, Y., Inoue, D., Ike, M. & Futamata, H. Learning beyond-pairwise interactions enables the bottom-up prediction of microbial community structure. Proc Natl Acad Sci U S A 121, e2312396121 (2024). 10.1073/pnas.2312396121

19 Diaz-Colunga, J., Catalan, P., San Roman, M., Arrabal, A. & Sanchez, A. Full factorial construction of synthetic microbial communities. bioRxiv, 2024.2005.2003.592148 (2024). 10.1101/2024.05.03.592148

20 Faust, K. et al. Microbial co-occurrence relationships in the human microbiome. PLoS Comput Biol 8, e1002606 (2012). 10.1371/journal.pcbi.1002606

21 Schafer, M. et al. Metabolic interaction models recapitulate leaf microbiota ecology. Science 381, eadf5121 (2023). 10.1126/science.adf5121

22 Maier, L. et al. Extensive impact of non-antibiotic drugs on human gut bacteria. Nature 555, 623–628 (2018). 10.1038/nature25979

23 Tramontano, M. et al. Nutritional preferences of human gut bacteria reveal their metabolic idiosyncrasies. Nat Microbiol 3, 514–522 (2018). 10.1038/s41564-018-0123-9

24 Hughes, C. V., Kolenbrander, P. E., Andersen, R. N. & Moore, L. V. Coaggregation properties of human oral Veillonella spp.: relationship to colonization site and oral ecology. Appl Environ Microbiol 54, 1957–1963 (1988). 10.1128/aem.54.8.1957-1963.1988

25 Dorison, L. et al. Identification of Veillonella parvula and Streptococcus gordonii adhesins mediating co-aggregation and its impact on physiology and mixed biofilm structure. mBio 15, e0217124 (2024). 10.1128/mbio.02171-24

26 Obana, N., Nakao, R., Nagayama, K., Nakamura, K., Senpuku, H. & Nomura, N. Immunoactive Clostridial Membrane Vesicle Production Is Regulated by a Sporulation Factor. Infect Immun 85 (2017). 10.1128/IAI.00096-17

27 Nakao, R. et al. A novel approach for purification and selective capture of membrane vesicles of the periodontopathic bacterium, Porphyromonas gingivalis: membrane vesicles bind to magnetic beads coated with epoxy groups in a noncovalent, species-specific manner. PLoS One 9, e95137 (2014). 10.1371/journal.pone.0095137

28 Jiang, Y., Kong, Q., Roland, K. L. & Curtiss, R., 3rd. Membrane vesicles of Clostridium perfringens type A strains induce innate and adaptive immunity. Int J Med Microbiol 304, 431–443 (2014). 10.1016/j.ijmm.2014.02.006

29 Boekhorst, J., de Been, M. W., Kleerebezem, M. & Siezen, R. J. Genome-wide detection and analysis of cell wall-bound proteins with LPxTG-like sorting motifs. J Bacteriol 187, 4928–4934 (2005). 10.1128/JB.187.14.4928-4934.2005

30 Siegel, S. D., Reardon, M. E. & Ton-That, H. Anchoring of LPXTG-Like Proteins to the Gram-Positive Cell Wall Envelope. Curr Top Microbiol Immunol 404, 159–175 (2017). 10.1007/82_2016_8

31 Momeni, B., Xie, L. & Shou, W. Lotka-Volterra pairwise modeling fails to capture diverse pairwise microbial interactions. Elife 6 (2017). 10.7554/eLife.25051

32 Stubbusch, A. K. M. et al. Antagonism as a foraging strategy in microbial communities. Science 388, 1214–1217 (2025). 10.1126/science.adr8286

33 Diaz-Colunga, J., Skwara, A., Vila, J. C. C., Bajic, D. & Sanchez, A. Global epistasis and the emergence of function in microbial consortia. Cell 187, 3108–3119 e3130 (2024). 10.1016/j.cell.2024.04.016

34 Palmer, J. D. & Foster, K. R. Bacterial species rarely work together. Science 376, 581–582 (2022). 10.1126/science.abn5093

35 Brinck, J. E. et al. Intestinal pH: a major driver of human gut microbiota composition and metabolism. Nat Rev Gastroenterol Hepatol 22, 639–656 (2025). 10.1038/s41575-025-01092-6

36 Ratzke, C. & Gore, J. Modifying and reacting to the environmental pH can drive bacterial interactions. PLoS Biol 16, e2004248 (2018). 10.1371/journal.pbio.2004248

37 Horvat, S., Mahnic, A., Breskvar, M., Dzeroski, S. & Rupnik, M. Evaluating the effect of Clostridium difficile conditioned medium on fecal microbiota community structure. Sci Rep 7, 16448 (2017). 10.1038/s41598-017-15434-1

38 Klieve, A. V., Yokoyama, M. T., Forster, R. J., Ouwerkerk, D., Bain, P. A. & Mawhinney, E. L. Naturally occurring DNA transfer system associated with membrane vesicles in cellulolytic Ruminococcus spp. of ruminal origin. Appl Environ Microbiol 71, 4248–4253 (2005). 10.1128/AEM.71.8.4248-4253.2005

39 Crost, E. H., Ajandouz, E. H., Villard, C., Geraert, P. A., Puigserver, A. & Fons, M. Ruminococcin C, a new anti-Clostridium perfringens bacteriocin produced in the gut by the commensal bacterium Ruminococcus gnavus E1. Biochimie 93, 1487–1494 (2011). 10.1016/j.biochi.2011.05.001

40 Chiumento, S. et al. Ruminococcin C, a promising antibiotic produced by a human gut symbiont. Sci Adv 5, eaaw9969 (2019). 10.1126/sciadv.aaw9969

41 Zhang, S. M., Hung, J. H., Yen, T. N. & Huang, S. L. Mutualistic interactions of lactate-producing lactobacilli and lactate-utilizing Veillonella dispar: Lactate and glutamate cross-feeding for the enhanced growth and short-chain fatty acid production. Microb Biotechnol 17, e14484 (2024). 10.1111/1751-7915.14484

42 Velsko, I. M., Chakraborty, B., Nascimento, M. M., Burne, R. A. & Richards, V. P. Species Designations Belie Phenotypic and Genotypic Heterogeneity in Oral Streptococci. mSystems 3 (2018). 10.1128/mSystems.00158-18

43 Hughes, C. S., Foehr, S., Garfield, D. A., Furlong, E. E., Steinmetz, L. M. & Krijgsveld, J. Ultrasensitive proteome analysis using paramagnetic bead technology. Mol Syst Biol 10, 757 (2014). 10.15252/msb.20145625

44 Mateus, A. et al. The functional proteome landscape of Escherichia coli. Nature 588, 473–478 (2020). 10.1038/s41586-020-3002-5

45 Kong, A. T., Leprevost, F. V., Avtonomov, D. M., Mellacheruvu, D. & Nesvizhskii, A. I. MSFragger: ultrafast and comprehensive peptide identification in mass spectrometry-based proteomics. Nat Methods 14, 513–520 (2017). 10.1038/nmeth.4256

46 Ritchie, M. E. et al. limma powers differential expression analyses for RNA-sequencing and microarray studies. Nucleic Acids Res 43, e47 (2015). 10.1093/nar/gkv007

